# Most sleep does not serve a vital function. Evidence from *Drosophila melanogaster*

**DOI:** 10.1101/361667

**Authors:** Quentin Geissmann, Esteban J. Beckwith, Giorgio F. Gilestro

## Abstract

Sleep appears to be a universally conserved phenomenon among the animal kingdom but whether this striking evolutionary conservation underlies a basic vital function is still an open question. Using novel technologies, we conducted an unprecedentedly detailed high-throughput analysis of sleep in the fruit fly *Drosophila melanogaster*, coupled with a life-long chronic and specific sleep restriction. Our results show that some wild-type flies are virtually sleepless in baseline conditions and that complete, forced sleep restriction is not necessarily a lethal treatment in wild-type *Drosophila melanogaster*. We also show that circadian drive, and not homeostatic regulation, is the main contributor to sleep pressure in flies. We propose a three-partite model framework of sleep function, according to which, total sleep accounts for three components: a vital component, a useful component, and an accessory component.

## Introduction

It is widely speculated that sleep serves a fundamental biological need, an idea derived from three distinct lines of evidence: (I) sleep is a universally conserved phenomenon across evolution; (II) chronic sleep restriction is often associated to death; (III) a sleepless animal has never been found (all reviewed in (1–4)).

The first of these three aspects – the striking evolutionary conservation of sleep – constitutes an important conundrum for scientists, but alone cannot be taken as a proof that sleep plays a vital function. Circadian rhythms, for instance, are also universally conserved, ultimately providing a clear evolutionary advantage, but they are not intrinsically vital to the individual given that animals can survive without a functional internal clock (5).

The fundamental question therefore is “can an animal survive without sleep?”. The study of chronic sleep deprivation could, at least in principle, address this challenge. Unfortunately, the literature on the chronic effects of sleep restriction is not comprehensive, partly dated and intrinsically complicated by the many confounding factors that correlate with sleep restriction. To date, experiments addressing this question have been reported in a handful of species only: dogs (reviewed in (6)), rats (reviewed in (7)), cockroaches (8), pigeons (9) and fruit flies (10). In four out of the five tested animal species, sleep deprivation experiments eventually terminated with the premature death of the animals but the underlying cause of lethality still remains unknown. In rats and dogs pups, death is associated with a severe systemic syndrome bearing important metabolic changes and clear signs of suffering, making it difficult to ultimately conclude whether lethality is caused by the mere removal of sleep or rather by the very invasive and stressful procedures employed to keep the animals awake (6, 11, 12). In the cockroach *Diploptera punctata*, sleep deprivation was achieved by continuously startling the animals (8), without however accounting for exhaustion-induced stress, a known lethal factor for other species of cockroaches (13–15). The observations in *Drosophila* are limited by the small number of animals used for the only reported experiment (12 flies in total) and by the methodology employed (human experimenters tapping their fingers on a tube (8)). In pigeons, chronic sleep deprivation was shown not to be lethal (9). In conclusion, chronic sleep deprivation experiments appear suggestive but inconclusive, for multiple reasons.

The third line of evidence supporting the hypothesis that sleep serves a fundamental biological need is perhaps the strongest and concerns the fact that sleepless individuals could never be identified, neither in nature nor through artificial laboratory screenings. We know that some species – such as elephants (16) or giraffes (17) – have evolved to cope with limited amount of sleep and several genetic mutations conferring short sleeping phenotypes in flies, rodents, and humans have been characterised in the past two decades (reviewed in (18)); some animals are also able to forego sleep for days or weeks in particular ecological conditions (16, 19–21), but the identification of a constantly sleepless animal can be considered a holy grail of the field.

Given that we do not possess a description of sleep at the cell-biological level, in all animals sleep quantification rely exclusively on *bona fide* macroscopic correlates, either electrophysiological or behavioural. Therefore, a technological development able to improve the characterisation of such correlates may provide a more accurate description of sleep, laying the conditions for a more specific sleep deprivation procedure. To this end, we recently created a system that allows for a faithful high-throughput analysis and manipulation of *Drosophila* sleep using activity as its behavioural correlate (ethoscopes (22)). Here, we report two surprising findings that were uncovered using such system, challenging the notion that sleep is a vital necessity: the discovery of virtually sleepless flies, and the finding that chronic sleep restriction is largely not lethal in *Drosophila melanogaster*.

## Results

### Virtually sleepless flies are found in a non-mutant population

Prolonged periods of inactivity are an evolutionary conserved, experimentally convenient behavioural correlate of sleep (23). Absence of movement is therefore routinely used as a proxy to measure sleep across a wide range of animals, spanning from jellyfish to elephants (16, 24, 25). In *Drosophila* too, sleep can be estimated by measuring the absence of walking bouts, generally using a commercially available device to detect whenever an isolated fly crosses the midline of a tube (26). This system, however, provides only limited spatial resolution which – unsurprisingly – results in an overestimation of sleep amounts (27). A growing number of laboratories are therefore transitioning to more accurate systems based on computer assisted video-tracking (27–31). To further improve our confidence in sleep estimation, we recently introduced a machine-learning approach that uses supervised learning to detect not only walking activity, but micro-movements too: *e.g*. in-place movements such as grooming, egg-laying, and feeding (22). How much do flies really sleep when, beside their walking activity, we measure their micro-movements too? To answer this question we analysed sleep for four consecutive days in 485 male (Fig. 1A) and 881 female (Fig. 1B) socially naive CantonS flies, a commonly used laboratory “wild-type” strain. As expected, in both males and females, sleep amounts were widely distributed, with male flies sleeping for 618.5, CI_95%_ = [606.7, 630.3], minutes a day and female flies for 299.2, CI_95%_ = [288.8, 309.6], minutes a day (mean, 95% bootstrap confidence interval; Fig. 1B). Interestingly, the distribution of sleep amount in females was wider than in males, with a long tail uncovering a previously undescribed fraction of short-sleeping female flies: 50% of female flies slept less than 20% of their time and 6% slept for less than 5% of their time (72 minutes a day). At the very end of the curve lied three flies that spontaneously slept an average of 15, 14, and 4 minutes a day respectively (Fig. 1A and Fig. S1). In both males and females, sleep amount is an endogenous feature: when flies are transferred into a novel environment – *i.e*. a fresh tube in a novel ethoscope inside a different incubator – their sleep amount remains mostly similar to their past sleep (Fig. 1C, R^2^ = 0.77, CI_95%_ = [0.73, 0.81]).

**Figure 1.**
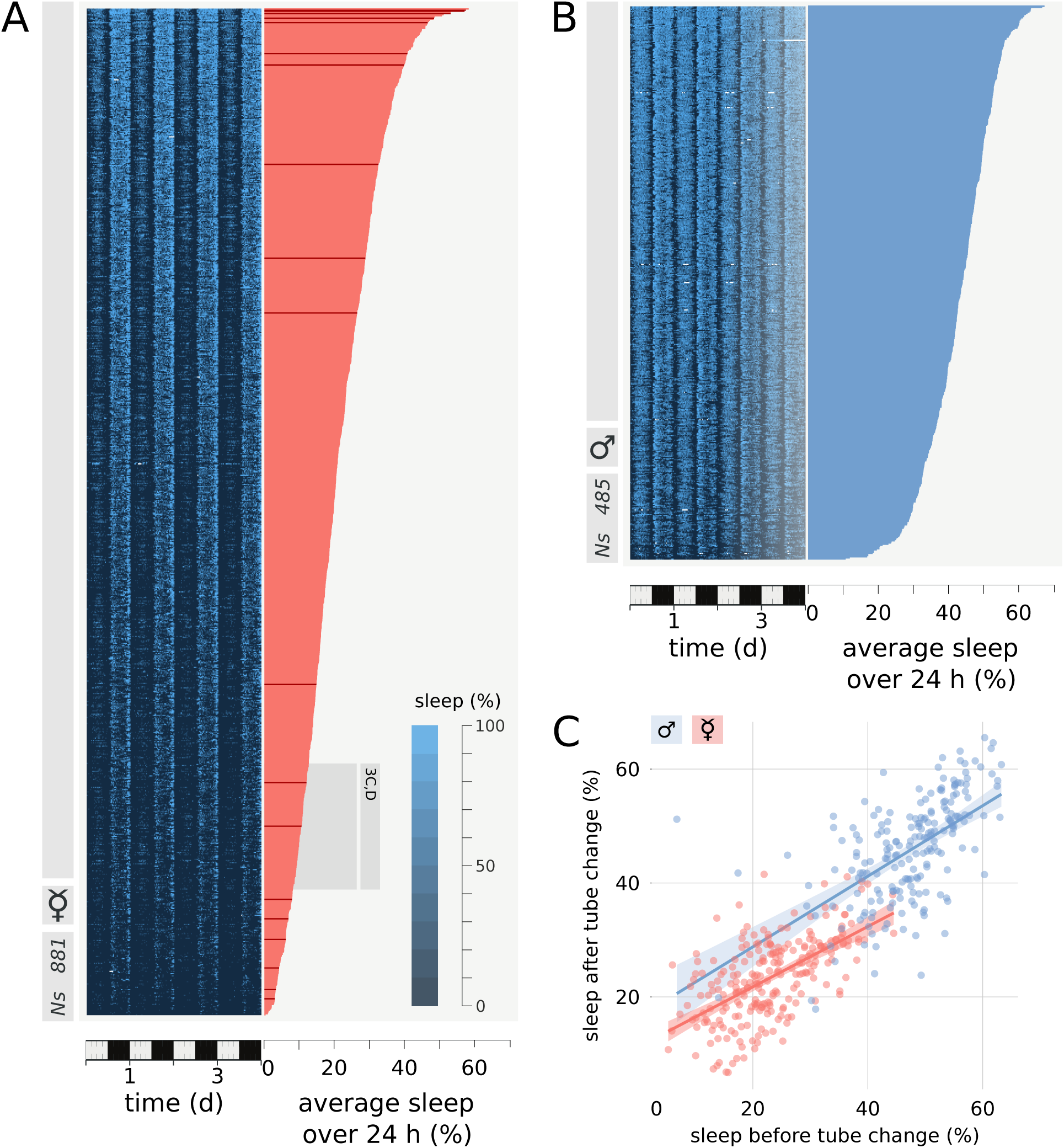
Great variability in sleep amounts in a non mutant population of *Drosophila melanogaster*. (A) Descending sorted distribution of sleep amount in a group of 881 female and (B) in a group of 485 male CantonS flies. The left panel shows sleep amount for each individual over a period of 5 days in bouts of 30 minutes. The right panel indicate the average sleep amount in 24 hours for female (pink in A and the rest of the figures) and male (cyan in B and the rest of the figures) flies. The group of short sleeping flies highlighted in grey is used for cluster analysis in Figure 3C and D. (C) Average sleep amount measured in a tube, predicts sleep amount measured in a different tube. Average of six days for both, with one day in between, N_male_ = 242 and N_female_ = 242.

### Micro-movements explain the short sleeping phenotype

Short sleeping flies have been identified in the past, either through experimental selection (32, 33), or through selected mutagenesis (34), but flies (and in fact animals) sleeping as little as few minutes a day where never identified before. To confirm the validity of our results, we reviewed the positional tracings of all 881 female flies in the dataset (Fig. S1), and acquired and reviewed videos for 19 flies with representative sleep amounts ranging from 823 to 42 minutes a day, to compare the tracking record at the single fly level (Fig. S2 – raw videos available at (35)). Manual inspection (Fig. S1, S2) and quantitative analysis (Fig. 2) confirmed that the activity repertoire oscillates in a stereotyped, sexually dimorphic, manner (Fig. 2A), with micro-movements being mostly present in females (Fig. 2B – in females, 623.4, CI_95%_ = [615.3, 631.4], minutes a day had at least one micro-movement episode, whilst in males 411.4, CI_95%_ = [404.2, 418.8], minutes did; mean, 95% bootstrap confidence interval). As expected, micro-movements (Fig. 2B) and movements that do not span the entire tube length (Fig. S1) are responsible for the quantitative difference in sleep analysis between recording platforms (Fig. 2C). Importantly, female micro-movements and quiescence are spatially (Fig. 2D) and, to a certain extent, temporally (Fig. 2A) exclusive: 37.3%, CI_95%_ = [36.9, 37.7], of the micro-movements happen at night (ZT12-ZT21), of which 51.3%, CI_95%_ = [50.3, 52.2], within 4 mm from the food (Fig. 2A and 2D respectively, green). In short, expanding on previously reported findings (36, 37), micro-movements in females are concentrated to those times of the day when flies are known to increase feeding activity (*i.e*. during mid-day and in the early phase of the night), mostly located by the food and away from the preferred site for quiescence (Movie S1 and Fig. 2), suggesting that the micro-movements observed in female flies are not a sleeping-related behaviour but a feeding-related behaviour.

**Figure 2.**
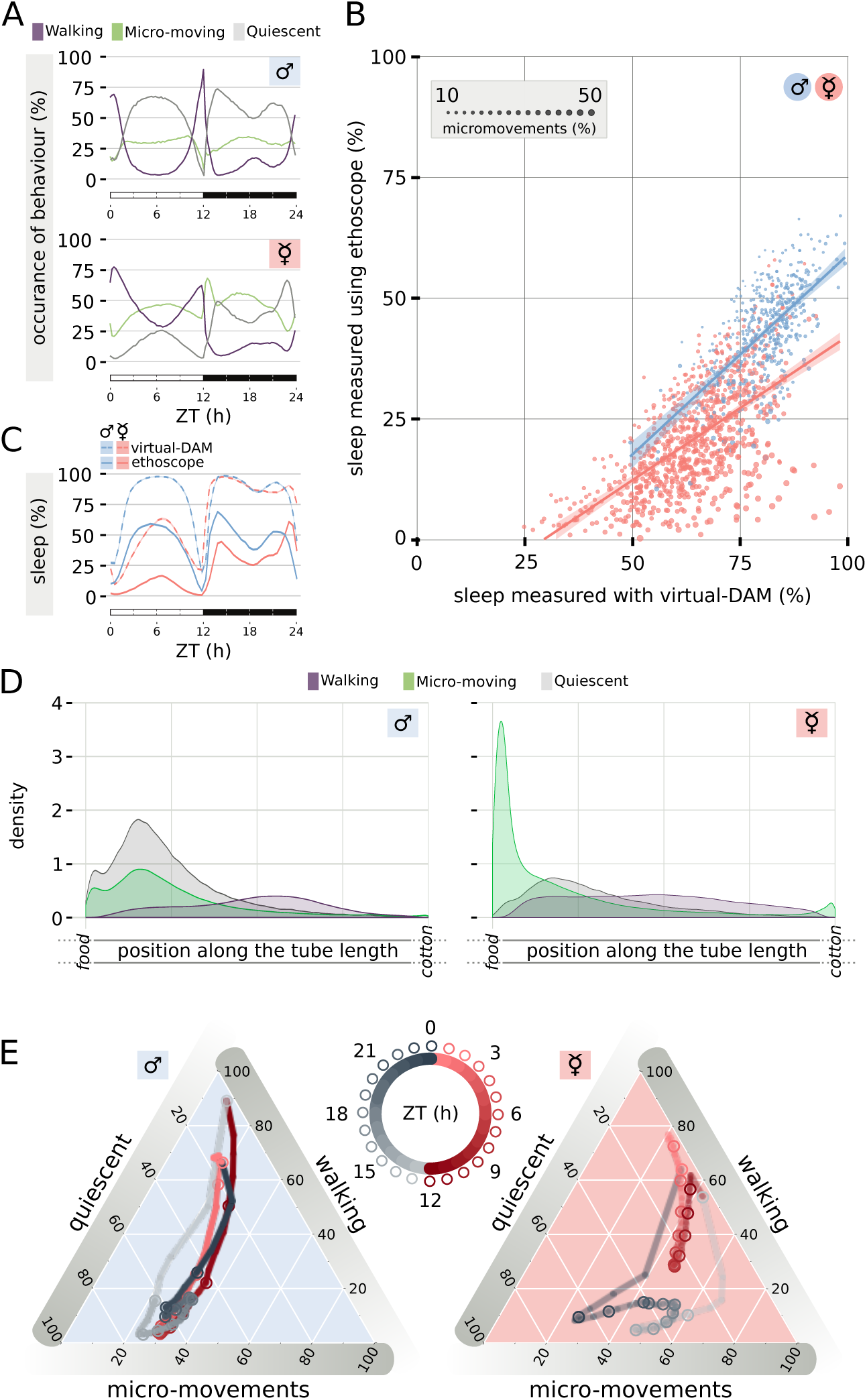
Micro-movements account for the newly described short sleeping phenotype. (A) Average occurrence of behaviour over the 24 h period in male (upper panel) and female (lower panel) Canton S flies. (B) Sleep amount for each individual male (cyan) and female (pink) fly plotted as computed with ethoscopes (y axis) and with virtual DAM analysis (29)(x axis). The size of each dots represents the average amount of micro-movements observed over the 24 hour period. (C) Average sleep amount over the 24 h period in male and female flies, plotted as computed with ethoscopes (continuous lines) or virtual DAM analysis (dashed lines). (D) Average positional distribution of behaviours for male (left panel) and female (right panel) flies over the 24 hour period, broken into the three behavioural states identified by ethoscopes. (E) Four dimensional representation of behavioural transitions over the 24 h period. Grey shades indicate the dark period (ZT 12-24), while red shades indicate the light period (ZT0-12). Same data set shown in Figure 1A,B.

### Qualitatively different types of short sleeping females

High-throughput ethoscope analysis allowed us to identify wild-type female flies that sleep as little as few minutes a day (Fig. 1B, Fig. S1). Could this be a peculiarity of some virgin flies, hence an ethological laboratory artefact? In *Drosophila*, mating status is known to be acting as a major behavioural switch (38) that modifies, among other behaviours, the animals dietary preference (36, 39) and their preference for feeding time (40). However, reaching sexual maturity only few hours after ecdysis, virgin female flies are likely to be a rare occurrence in the wild (41, 42). To test how sleep changes with mating status, we recorded sleep in flies before and after a successful (green), or unsuccessful (grey), mating event (Fig. 3A). In mated females, sleep dramatically decreased for at least three consecutive days (Fig. 3A – in the first full day after mating, lowering from 383.2, CI_95%_ = [363.4, 404.9], to 175.3, CI_95%_ = [153.0, 200.2] minutes a day), correlating with a major change of positional preference of the animals towards the food (Fig. 3B). Such change in positional preference (Fig. 3B) and the strong increase in micro-movements (Fig. 3C) are likely to represent an increase in food intake and egg-laying activity and may explain why such a strong decline in sleep amount was never identified using different tools (27, 43, 44). Interestingly, four-dimensional behavioural fingerprinting showed that the short sleep phenotype observed upon mating is qualitatively different from the one observed as natural variation in the CantonS population (Fig. 3C, D and Fig. S2).

**Figure 3.**
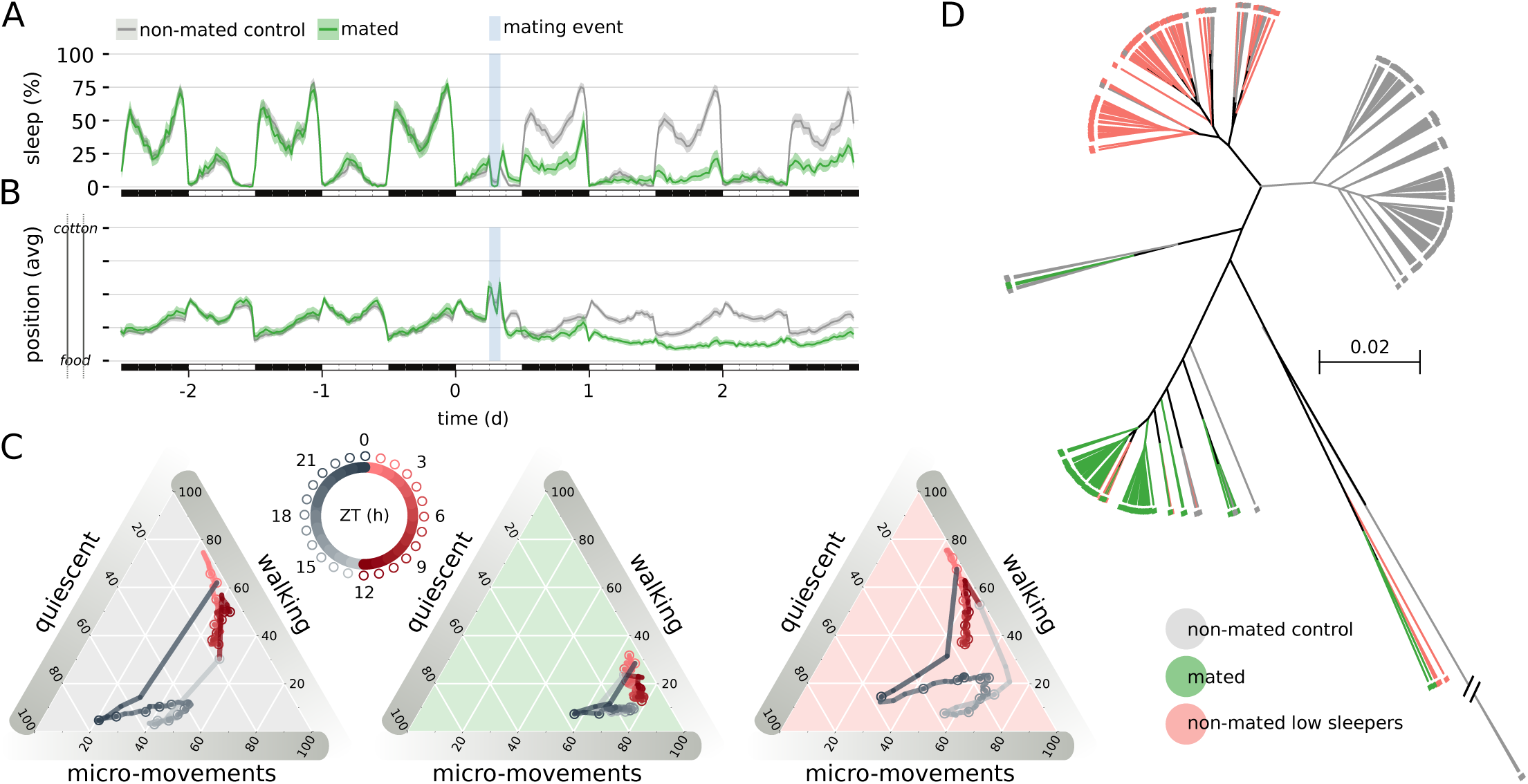
Mating reduces sleep amount. (A) Sleep profile of all the female flies used in the mating experiment: Green: flies that underwent successful mating event (N = 86); Grey: flies that met a male but did not engage in copulation (N = 152). The blue vertical shade indicates the timing of the mating event. (B) Average position along the tube of the same flies shown in A in 30 minutes bins. (C) Four dimensional representation of behavioural transitions over the 24 h period for non-mated flies (grey background), mated flies (green background) and for naturally short sleeping unmated flies (pink background – same dataset highlighted in grey in Figure 1A). (D) Hierarchical clustering based on pairwise distance, in the time-behaviour domain, of the same three cohorts shown in C.

### Prolonged sleep deprivation is largely non lethal

The experiments described so far unveiled that a fraction of female flies necessitate little sleep, with some being almost completely sleepless. Is sleeplessness a peculiarity of a few special individuals, or can any fruit fly cope with little or no sleep? To answer this question, we conducted a life-long sleep deprivation experiment using a closed-loop sleep deprivation device able to interact with single animals by triggering a tube rotation after a predefined period of immobility (22), a system created to minimize the extent of disturbance and conceptually inspired by the disc-overwater apparatus developed by the Rechtschaffen laboratory (7) in which a rat receives a waking physical challenge only when it is factually asleep, but is left undisturbed otherwise. In our setup, flies were housed in individual tubes and each tube experienced a 1 second rotation at the approximate speed of 300 rpm whenever the animal housed inside had shown 20 s of continuous immobility (*22*). The treatment led to a highly efficient sleep deprivation with flies losing, on average, 95.6%, CI_95%_ = [93.5 98.2], of their sleep (Fig. 4A) and yet, surprisingly, we could not detect any major effect on survival (Fig. 4B, C). In particular, sleep deprived male flies lived as long as the control group (with a median of 41.5, CI_95%_ = [38.0, 44.0], days against 46.0, CI_95%_ = [41.0, 48.5], days for the controls) and a minor effect was only evident in female flies, with a reduction of median lifespan of 3.5 days (37.5, CI_95%_ = [33.0, 38.5], 41.0, CI_95%_ = [38.5, 44.0]). Forced sleep restriction is largely not lethal in flies when performed in a controlled, specific, manner.

**Figure 4.**
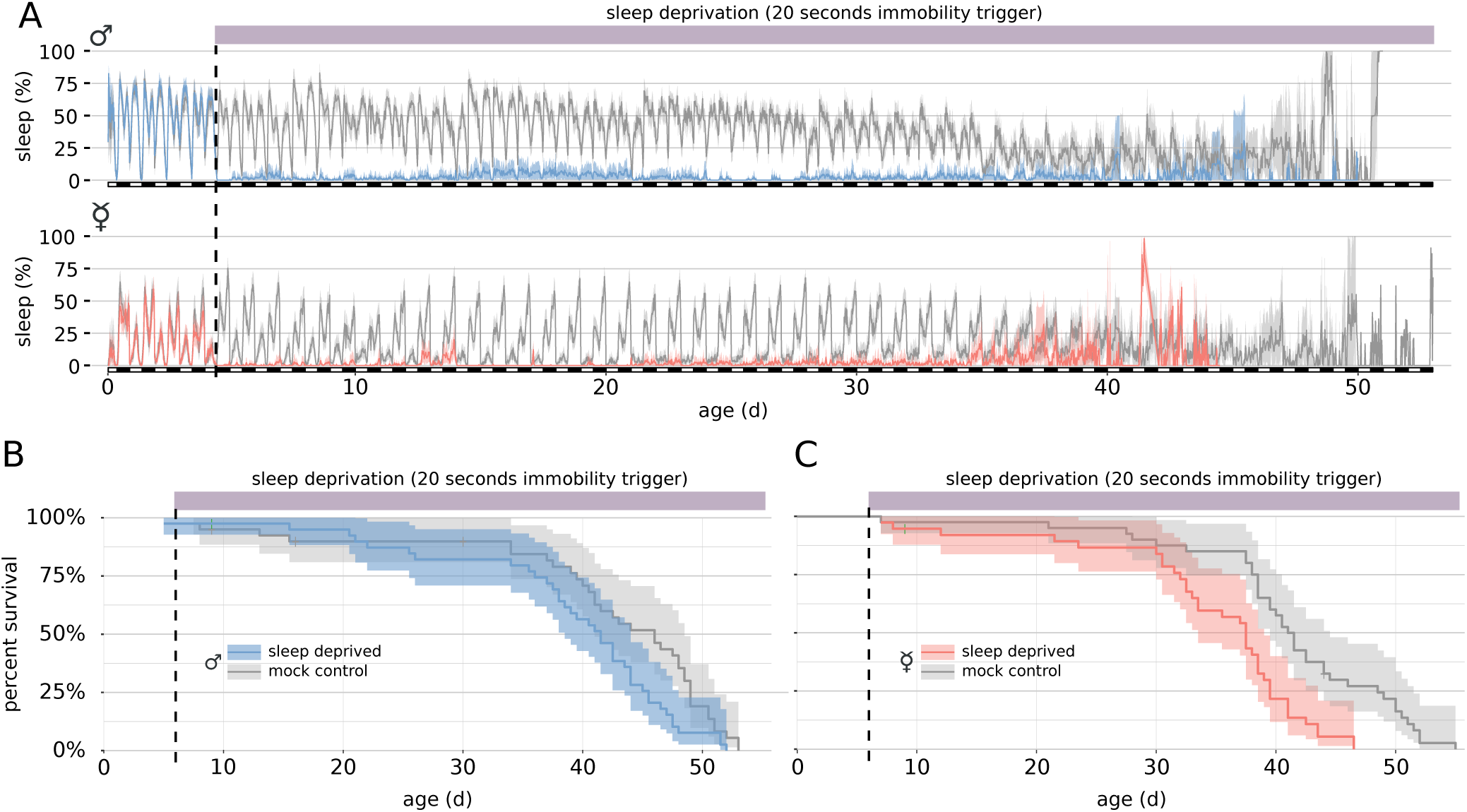
Chronic mechanical sleep deprivation is largely not lethal in *Drosophila melanogaster*. (A) Life-long sleep restriction in male (upper panel) or female (lower panel) CantonS flies subjected to mechanical sleep deprivation triggered by a 20 s inactivity bout. (B) Survival curve for male (cyan in B) or female (pink in C) sleep deprived flies and their sex-matched undisturbed control (grey in both B and C). Sleep measurements became noisier as the number of flies decreases. N = [38,40] for all four groups.

### Sleep rebound after sleep deprivation only partly correlate with sleep loss

If most (all?) sleep does not serve a direct and immediate vital function, do we need to rethink the current prevailing concept of sleep homeostasis? Is sleep rebound a way to make up for a loss of an otherwise impaired biological process, or is it instead merely a “punishment” phenomenon, evolved to guarantee that a constant, largely species-specific amount of sleep is met? To explore this new dichotomy, we analysed how different treatments of sleep deprivation would affect sleep rebound. To start, we conducted an acute sleep deprivation experiment on a total of 818 male (Fig. 5A-D) and 992 female (Fig. 5E-H) CantonS flies, with a comprehensive range of immobility triggers, spanning from 20 to 1000 seconds, to deprive flies of sleep episodes of specific length. As expected, the total amount of sleep lost during the 12 h of deprivation positively correlated with the length of the immobility trigger adopted (Fig. 4B,F) whilst the number of stimuli delivered was inversely correlated (Fig. 4C,G). Interestingly, in all cases could we observe a statistical significant sleep rebound in the first 3 hours following the sleep deprivation, also when the sleep loss was not statistically different than control (Fig. 4F, 840 s and 1000 s inactivity triggers). In particular, depriving female flies of only the longest sleep episodes (≥1000 s) still led to a significant sleep rebound the subsequent morning, even though flies experienced, on average, only 5.8, CI_95%_ = [4.8, 6.8], of tube rotations per night.

**Figure 5.**
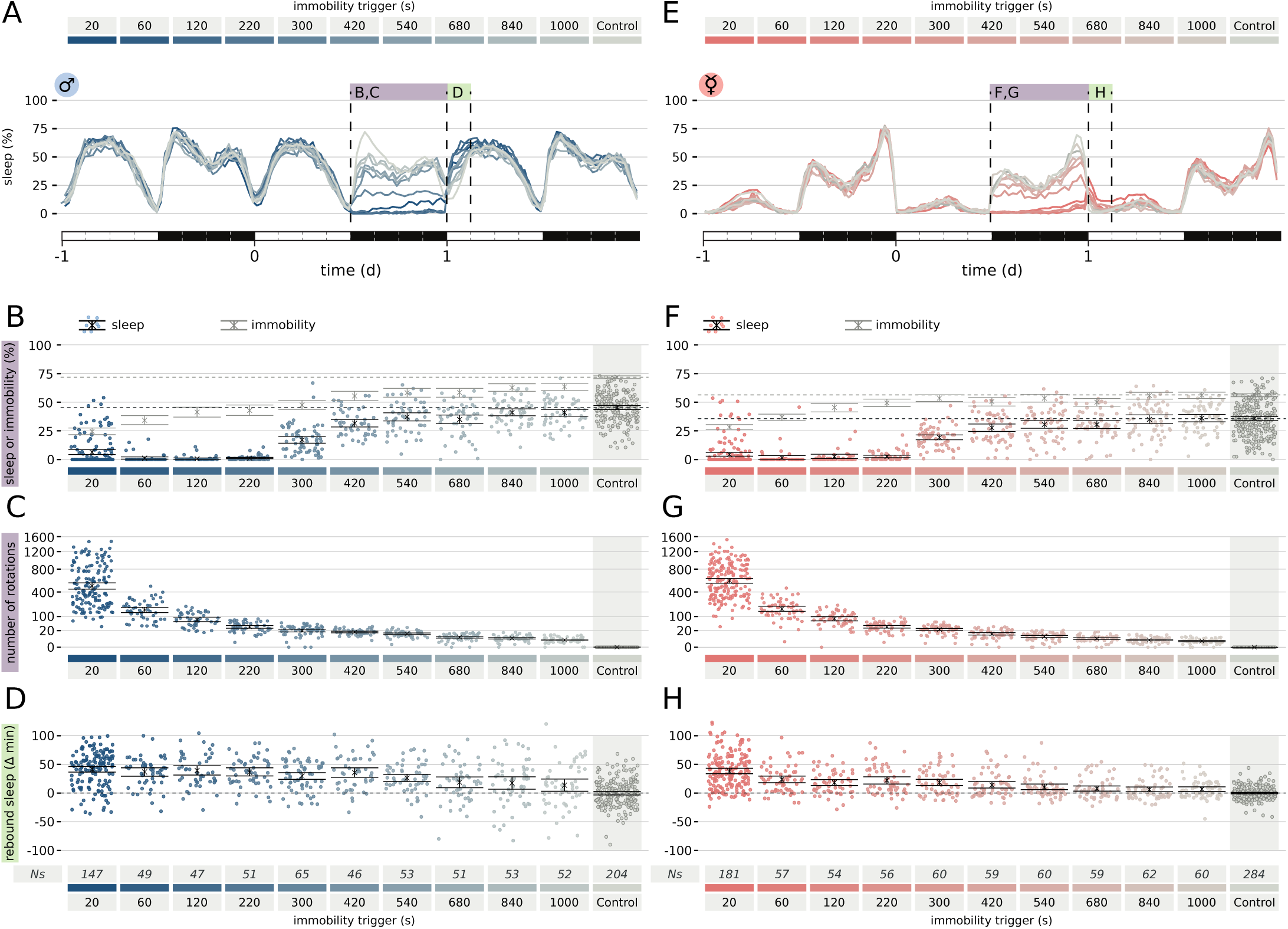
Sleep rebound is not linearly proportional to sleep loss. (A, E) Sleep profile for the entire data set shown: 818 male (A-D) and 912 female (E-H) CantonS flies. (B, F) Sleep (cyan and pink dots and black lines) or immobility (grey lines) for the entire data set spanning 10 different immobility interval triggers (20 s to 1000 s). Control flies were never actively stimulated but laid adjacently to the experimental flies. (C, G) Number of tubes rotations triggered by immobility bouts. (D, H) Amount of rebound sleep in the ZT0-3 interval following the sleep deprivation for the entire data set.

The increase in sleep pressure driving rebound after sleep deprivation is not linearly correlated with the amount of sleep lost over the length of one night, but how do flies react to prolonged sleep restriction spanning multiple days? To answer this question, we conducted a “Randy Gardner”-like experiment (45), in which we subjected flies to 228 h of uninterrupted sleep deprivation, using a 20 s immobility trigger as waking event. The experiment was conducted both in males and females flies, using undisturbed control individuals in adjacent tubes, for a total of 377 animals (Fig. 5). Even after almost 10 d of chronic sleep deprivation, male flies manifested a sleep rebound that was not dissimilar from the rebound observed after one night of acute sleep restriction (visually compare Fig. 6C to Fig. 5A). Intriguingly, whilst in male flies rebound sleep was again limited to the first three hours of rebound day, in female flies the observed sleep rebound was quantitatively modest but appeared to be protracted in time for the subsequent three days, at least (Fig. 6B and 6D). Because the tube rotations were triggered by immobility, we could use the number of rotations (Fig. 6F, dashed lines) and the distance walked (Fig. 6F, continuous lines) as proxy of endogenous sleep pressure in control (grey) or sleep deprived (pink and blue) flies (Fig. 6F).

**Figure 6.**
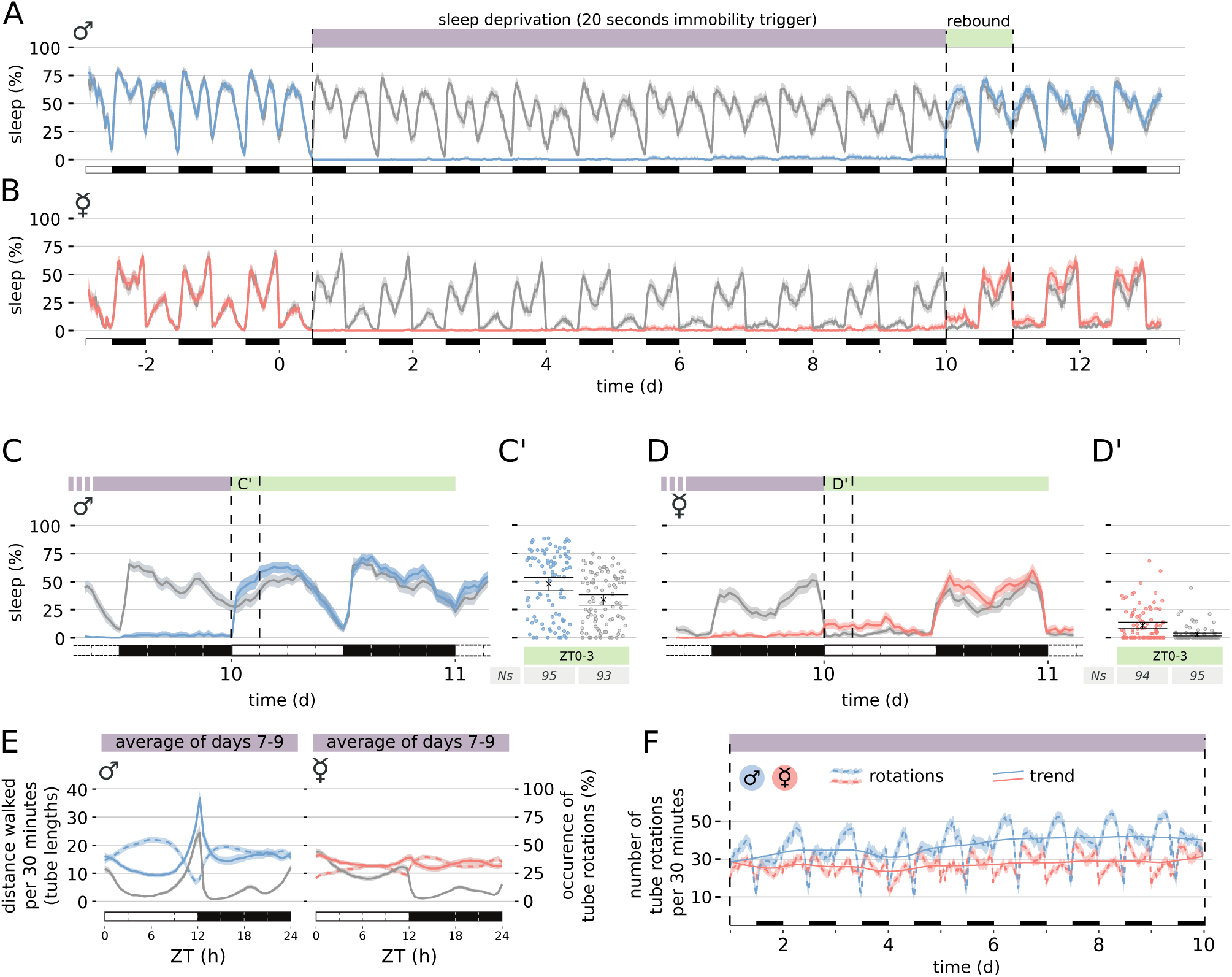
Sleep pressure is largely under control of the circadian rhythm. (A, B) Sleep profile for male (A, cyan) and female (B, pink) CantonS flies during the length of the experiment compared with their sex-matched undisturbed controls (grey in both). Day 0 signs the beginning of the chronic sleep deprivation procedure, lasting 228 hours (indicated by a purple shade on top). The green shade indicates the rebound day blown up in C and D. (C, D) Magnification of the sleep deprivation to rebound transition. (C’, D’) Quantification of sleep amount during ZT0-3 of rebound day. (E) Activity of flies shown as distance walked (continuous lines: grey, control; cyan, males; pink, females) or, by proxy, as number of rotations over the average 24 hours period (dashed lines: cyan, males; pink, females). (F) Actual average number of tubes rotations over the length of the sleep deprivation experiment (dashed lines) or seasonal trend (continuous lines – see supplementary methods for details). N = [93, 95] for all four groups.

In both male and female flies, the main changes in sleep pressure were cycling in a circadian fashion, with the clock-regulated bouts of walking activity still showing no sign of subsidence, despite the long sleep deprivation (Fig. 6F). In other words: when the circadian clock commands activity, the flies are active also after days and days of cumulating sleep pressure. In fact, seasonal decomposition of rotations over the 9.5 d of sleep deprivation confirmed that only a small amount of the variance in sleep pressure is explained by the long range trend in sleep deprivation (21% in males and 11% in females, Fig. 5F), whilst the main contributor of sleep pressure is indeed circadian periodicity (69% in males and 61% in females, Fig. 5F continuous lines). These data, taken together, clearly indicate that the main stimulus to rest in flies is driven by the circadian clock.

## Discussion

The idea that sleep fulfils a vital biological need – we initially argued – relies on one fundamental question: can we find an animal able to survive without sleep? According to the data presented here, the answer could be “ye”s. In wild type *Drosophila melanogaster* the need for sleep is not a vital necessity and lack of sleep – either endogenously driven (Fig. 1) or artificially imposed (Fig. 4 and 6) – is compatible with life. The utmost conceptual importance of these findings commands caution, and caveats must be critically examined. Most importantly, we cannot rule out that, in our experiments, flies still experience enough sleep to satisfy an hypothetical vital need. In other words, prolonged or consolidated sleep is not a vital necessity but intervals of sleep that last only few seconds (20 s in most experiments here presented) may be sufficient to satisfy whatever basic biological need sleep may serve. Behavioural correlates of sleep have been described in virtually every animal that has been studied so far, connecting species as different as jellyfishes and humans (25, 46), and have demonstrated that sleep amounts vary dramatically across the animal kingdom. For instance, elephants sleep as little as 3 hours a day (16), Tinaja cavefish as little as 2 hours a day (47), whereas little brown bats sleep, on average, 20 hours a day (46). No existing model of sleep function can account for this variability. One intriguing possibility, which we propose to the reader here, is that sleep should not be seen as a monolithic phenomenon but rather as the mixture of three components: a *vital* component, a *useful* component, and an *accessory* component. The accessory component is conceptually identical to what has been previously coined “adaptive inactivity” (2) or “trivial function of sleep” (3) and would postulate that at least a fraction of sleep would serve no core biological function other than circadianly syncing periods of wakefulness in the most ecologically appropriate manner, for instance keeping animals out of danger or restricting their activity to gain safety and rest. Can we accept that a good fraction of those 20 hours of sleep a brown bat requires is driven by the evolutionary adaption of staying “out of trouble”? And, if this is intuitively easy to accept for bats, why should it not be universally true? Other vital needs, such as feeding, follow a similar three-partite subdivision, with a given amount of calories and nutrients being vitally needed, some useful, and some merely accessory – and even detrimental. In our experiments, we may have removed the last two components (accessory and useful) but left enough sleep to satisfy a yet mysterious vital need, even if in the form of short bouts lasting few seconds each. We also uncovered an interesting sexual dimorphism in terms of undisturbed sleep need and in terms of response to sleep deprivation: whilst female flies are able to cope with much less sleep in baseline conditions, they are more sensitive to sleep deprivation, with an extended rebound upon long sleep restriction (Fig. 6B and D) and a moderate but significant effect on lethality upon life-long sleep deprivation (Fig. 4). This sexual dichotomy may be instrumental in the future to dissect the difference between the three-partite components.

At first sight, the results presented here appear to be clashing with some of the existing knowledge. In our view, they command, instead, for a thorough review of existing sleep deprivation literature. The experiments of chronic sleep deprivation performed in dogs pups at the end of 1800s are universally considered too primitive to be trustworthy and too unethically stressful to be reproducible in modern times (6). The early *Drosophila* experiments were too preliminary to depict a whole picture, marred by a limited number of animals (12 individuals) and by the adoption of a procedure that is not easily reproducible (human experimenters finger tapping on the tubes) (10). Other lines of research have also shown no correlation between sleep loss and survival in flies: loss of the *insomniac* (48) or *fumin* (34) genes leads to strong sleep restriction that is still compatible with life. Likewise, artificially selected short-sleeping fruit flies have unaltered longevity (32). With flies joining pigeons in the list of animals surviving chronic sleep deprivation, the only solid evidence in favour of lethality upon sleep deprivation lays with the chronic sleep deprivation in rats using the disc-over-water system. Those experiments, however, were not free of confounding factors and one cannot exclude a stress or metabolic component given that animals were thrown into water several hundreds times a day (12). In humans, for obvious ethical reasons, we have no experimental evidence that prolonged sleep deprivation is incompatible with life. A human prion diseases, fatal familial insomnia (FFI), is sometimes brought as evidence of a vital function of sleep, yet bearing too many confounding factors, considering the devastating nature of the pathology (49). Importantly, transmitted (50) and transgenic mouse models (51) of FFI reproduce clear signs of neurodegeneration and premature death, but not sleeplessness suggesting that, in humans, the insomnia is a symptom of the disease but not necessarily the cause of death (52). In conclusion, we believe our results clearly show that the time is ripe for the field to revisit the dogmatic believe that sleep serves a unique, evolutionary conserved, function.

## Acknowledgements

We thank all members of the Gilestro laboratory for endless discussions and invaluable intellectual contributions.

## Author-contributions

EJB and QG performed the experiments; all authors analysed the data and contributed to the manuscript.

## Data and material availability

all data, raw and processed, is available in the manuscript or the supplementary materials.

## Supplementary Materials

### Materials and Methods

**Figure S1.** Representative tracings of the behavioural activity over the course of 48 h as recorded in real-time by the ethoscopes for all 818 female flies shown in Figure 1A. The continuous black line plots the position of the flies along the tube (y axis) over time (x axis). The transversal dashed line represents the position of a virtual infrared beam (*29*). The colour on the background highlights the concomitant behavioural classification with a resolution of 1 minute (grey: quiescent, green: micro-moving; blue: walking).

**Figure S2.** Sorted hierarchical cluster analysis based on pairwise distance, as supplement to Figure 3. (A) Average daily sleep amount as % of the day for each female fly in the dataset. Same colour code as Figure 3. (B) Hierarchical clustering dendrogram based on pairwise distance. (C) Visual representation of the average occurrence of the three behavioural features in each single animal across the 24 hour period.

**Movie S1.** Visual representation of the distribution of behaviour features across the 24 h in the dataset shown in Figures 1A,B and 2. Each frame in the movie bins data with a 15 minutes resolution: red: female flies; blue: male flies. In the leftmost panels, each dot is an individual animal plotted in their behavioural space at that time point. The right most panel shows the 24 h distribution of the following four features: fraction of queiscent, micro-moving, walking animals (top three) and average position along the tube longitudinal length (bottom).

